# Pytometry: Flow and mass cytometry analytics in Python

**DOI:** 10.1101/2022.10.10.511546

**Authors:** Maren Büttner, Felix Hempel, Thomas Ryborz, Fabian J. Theis, Joachim L. Schultze

**Affiliations:** German Center for Neurodegenerative Diseases (DZNE), Bonn, Germany; Life and Medical Sciences (LIMES) Institute, University of Bonn, Germany; Institute of Computational Biology, Helmholtz Center Munich, Germany; University of Applied Sciences Weihenstephan-Triesdorf, Freising, Germany; TUM School of Life Sciences Weihenstephan, Technical University of Munich, Germany; PRECISE Platform for Single Cell Genomics and Epigenomics, Deutsches Zentrum für Neurodegenerative Erkrankungen (DZNE) and the University of Bonn, Bonn, Germany

## Abstract

Flow and mass cytometry data are commonly analyzed via manual gating strategies which requires prior knowledge, expertise and time. With increasingly complex experiments with many parameters and samples, traditional manual flow and mass cytometry data analysis becomes cumbersome if not inefficient. At the same time, computational tools developed for the analysis of single-cell RNA-sequencing data have made single cell genomics analysis highly efficient, yet they are mostly inaccessible for the analysis of flow and mass cytometry data due to different data formats, noise assumptions and scales. To bring the advantages of both fields together, we developed Pytometry as an extension to the popular scanpy framework for the analysis of flow and mass cytometry data. We showcase a standard analysis workflow on healthy human bone marrow data, illustrating the applicability of tools developed for the larger feature space of single cell genomics data. Pytometry combines joint analysis of multiple samples and advanced computational applications, ranging from automated pre-processing, cell type annotation and disease classification.

## Introduction

With the lack of computational automation, but also due to technical limitations of flow cytometry during the time of its introduction to the life sciences, data analysis relied on manual gating strategies, which required prior knowledge as well as expertise. As of today, manual gating is the current standard for analyzing flow cytometry data, despite it being well-known and accepted in the field that manual gating suffers from several shortcomings. The analysis process is manual, thus, different experts annotate the data differently, and one person may analyze the data differently over time. Moreover, manual gating is very difficult to scale to hundreds or thousands of samples - a number that is increasingly presented in clinical cohorts.

At the same time, gating usually targets populations of interest and one rarely performs exhaustive gating for all cell populations within the dataset. This becomes especially relevant when analyzing data generated with disease models: perturbed cell populations may be missed with common gating strategies due to their aberrant expression of markers. Further, panel sizes for flow cytometry data increased well above 20 markers in recent years, which renders exhaustive gating for all cell populations next to impossible and urges the use of computational analysis for instance through dimensionality reduction and clustering. Such approaches have been used to analyze mass cytometry data with typical panel sizes of about 40 markers (Bruggner *et al*., 2014; Qiu *et al*., 2011; Pezzotti *et al*., 2016). Notably, with the advent of multi-modal single-cell omics technologies such as CITEseq (Stoeckius *et al*., 2017) and Abseq (Shahi *et al*., 2017), the need to directly compare these data to flow or mass cytometry becomes imminent.

Finally, machine learning based analysis of flow data processes millions of measurements at a time and requires efficient handling of memory and computation time. We aim to unlock the potential of cytometry data for automated analysis and classification workflows. Here, we present Pytometry, a Scanpy (Wolf *et al*., 2018) extension to analyze flow and mass cytometry data in Python.

## Description: The Pytometry Python package

The Pytometry package extends on the single-cell analysis framework scanpy. Commonly, flow or mass cytometry data are stored in FCS file format. Pytometry converts FCS files into the annotated data frame format (Sun and Wolf, 2022) of the anndata package (Virshup *et al*., 2021) and stores the header information alongside with the data matrix. For each feature, Pytometry stores both the marker name and the corresponding color for flow cytometry data. Marker names may have several alternative annotations, which are stored in the metadata table of the features. The focus of the pytometry package is processing and comparing multiple samples with millions of cells at once, in contrast to flow analysis packages like FlowKit (White et al. 2021) and CytoPy (Burton et al. 2021).

For the analysis of flow cytometry data, one usually has to compensate for the fluorescent spillover. Pytometry uses the compensation matrix provided in the FCS file to perform the compensation. The user may also provide a custom compensation matrix. Next, Pytometry provides built-in flow cytometry specific data normalization, such as bi-exponential and logicle transformation (adapted from FlowKit (White *et al*., 2021) for the anndata format), and the inverse sinus hyperbolicus (arcsinh) transformation for mass cytometry data. The data are then further processed using the scanpy framework for low dimensional embedding such as UMAP (Becht *et al*., 2018; McInnes and Healy, 2018) and clustering using the Leiden community detection algorithm (Traag *et al*., 2019). Batch effect correction can be achieved with over 16 different single-cell tailored methods via the scIB package (Luecken *et al*., 2022). Finally, writing data to file is handled via storage solutions within the anndata package, like zarr and the language independent HDF5 file format.

## Application

We applied Pytometry to analyze a single-cell mass cytometry data set of eight healthy human bone marrow donors (Oetjen *et al*., 2018) with a panel of T-cell-specific markers (Figure 1). We loaded the data from the FCS file format into the anndata format and merged all data matrices. We then normalized using the arcsinh transformation with cofactor 5. Next, we selected only immune cells that are positive for CD45. From the remaining 4.1 million cells, we computed a neighborhood graph and a UMAP (Figure 1a-b). To annotate the subsets of T cells, we used Leiden clustering to group the cells and renamed the clusters according to the marker intensity of NK-cell and T-cell specific markers (Figure 1c, Supplementary Table 1 and Supplementary Material of (Oetjen *et al*., 2018)). In particular, scanpy allows hierarchical annotation of cell types from general cell types to highly resolved subtypes (Figure 1a) and to display the corresponding marker intensity (Figure 1c). Clustering recovers all cell types and compositions (Figure 1d-e). Cell type abundance from clustering was consistent with the expert analysis results of conventional flow and mass cytometry analysis (Oetjen *et al*., 2018).

**Figure 1.**
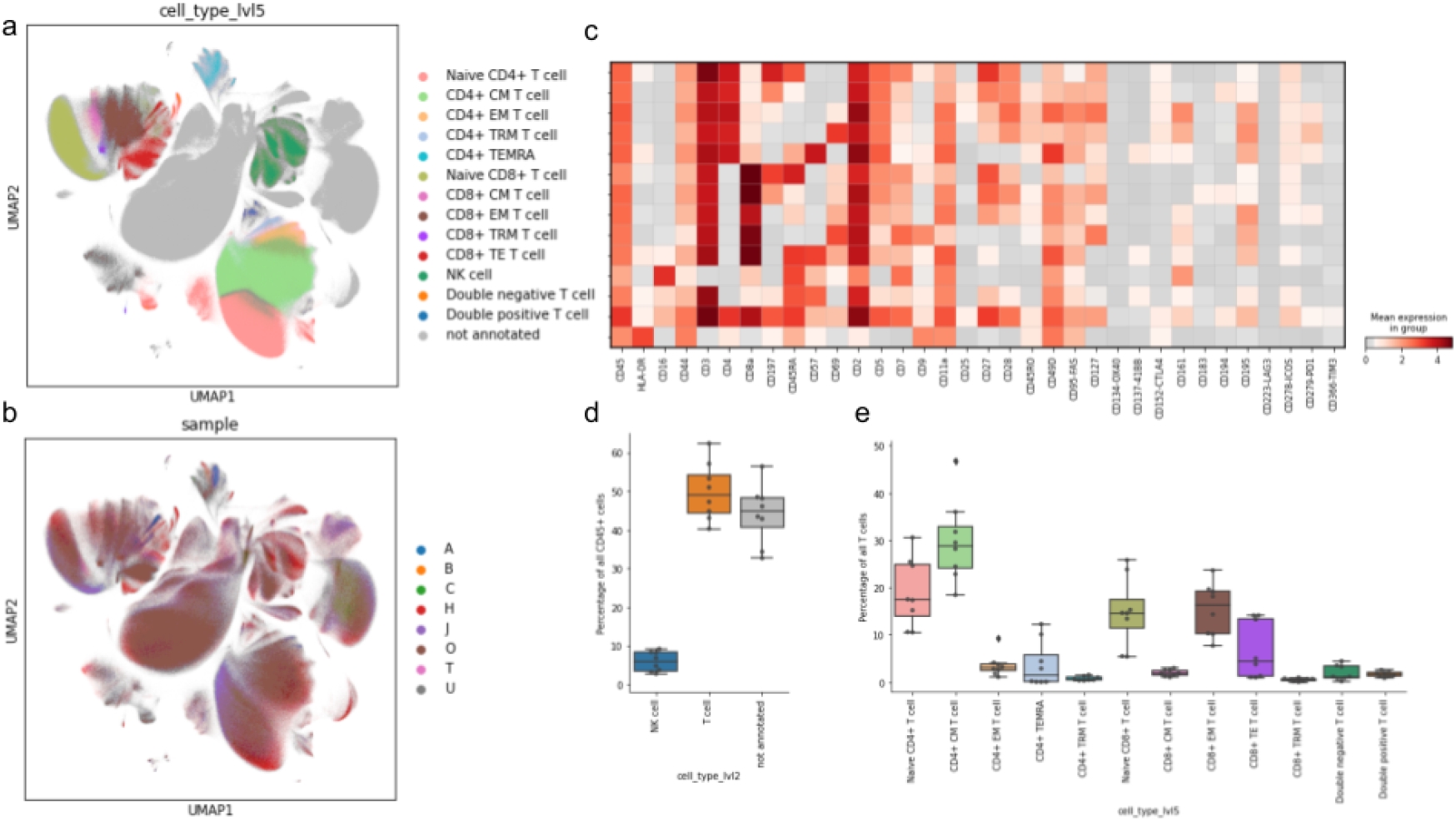
*Analysis of single-cell mass cytometry data from human bone marrow samples. (a) UMAP of all CD45+ cells. Gray cells are not annotated due to lack of distinctive markers. (b) UMAP colored by donor. (c) Matrixplot of the mean marker intensity per cell type. (d) Boxplot of NK and T cell type proportions per donor. Dots indicate the proportion of each donor. Boxes indicate 25th and 75th percentile. The horizontal bar in boxes indicates the median. Whiskers indicate the overall range of values. (e) Boxplot of T cell type proportions per donor. Dots indicate the proportion of each donor. Boxes indicate 25th and 75th percentile. The horizontal bar in boxes indicates the median. Whiskers indicate the overall range of values*.

## Discussion and conclusion

We presented a user-friendly Python package for the analysis of flow and mass cytometry data. While packages like Flowkit (White *et al*., 2021) and cytopy (Burton *et al*., 2021) build on their own data structure, Pytometry builds upon the annotated data frame (anndata) and scanpy packages, and allows efficient handling of flow and mass cytometry data in Python. It integrates fully with the scverse ecosystem and ensures long-term maintenance. We demonstrate its capabilities for analyzing 4.4M cells from mass cytometry experiments. Our package is able to detect all cell types and their abundance. Pytometry serves as the basis for a multitude of machine learning applications on flow data, ranging from automated pre-processing, cell type annotation and disease classification.

## Data availability

We downloaded the publicly available flow and mass cytometry data of healthy human bone marrow donors (Oetjen *et al*., 2018) from FlowRepository.org (accession codes: FR-FCM-ZYQ9, FR-FCM-ZYQB).

## Code availability

The Pytometry Python package is available on Github https://github.com/buettnerlab/pytometry.

## Acknowledgements

We thank Sunny Sun and Alex Wolf from Lamin Labs for their support on the code base. We acknowledge Lorenzo Bonaguro’s feedback on the mass cytometry data analysis. M.B. acknowledges funding by the iTreat-BMBF (DLR) (grant # 01ZX1902B). F.J.T. acknowledges funding by the Helmholtz Association’s Initiative and Networking Fund through Helmholtz AI (grant # ZT-I-PF-5-01).

## Author contributions

M.B., F.J.T. and J.L.S. designed the study, supervised the work and wrote the manuscript with the help of all co-authors. F.H., T.R. and M.B. wrote the code and performed the analysis.

## Supplementary Material

**Supplementary Table 1:** Marker combination for annotating NK cells and T cells

| Cell Type | Marker combination |
| --- | --- |
| NK cell | CD45+ CD16+ HLA-DR- CD3- CD44- CD45RA+ |
| T cell | CD45+ CD3+ |
| CD4+ T cell | CD45+ CD3+ CD4+ |
| CD8+ T cell | CD45+ CD3+ CD8a+ |
| Double positive T cell | CD45+ CD3+ CD4+ CD8a+ |
| Double negative T cell | CD45+ CD3+ CD4- CD8a- |
| Naive CD4 T cell | CD45+ CD3+ CD4+ CD197+ CD45RA+ |
| Naive CD8 T cell | CD45+ CD3+ CD8a+ CD197+ CD45RA+ |
| CD4+ Central Memory (CM) T cell | CD45+ CD3+ CD4+ CD197+ CD45RA- |
| CD8+ CM T cell | CD45+ CD3+ CD8a+ CD197+ CD45RA- |
| CD4+ Effector Memory (EM) T cell | CD45+ CD3+ CD4+ CD197- CD45RA- |
| CD8+ EM T cell | CD45+ CD3+ CD8a+ CD197- CD45RA- |
| CD4+ Tissue-resident T cell (TRM) | CD45+ CD3+ CD4+ CD197- CD45RA- CD69+ |
| CD8+ TRM | CD45+ CD3+ CD8a+ CD197- CD45RA- CD69+ |
| CD4+ terminally differentiated effector memory T cells (TEMRA) | CD45+ CD3+ CD4+ CD197- CD45RA+ |
| CD8+ Effector T cell (TE) | CD45+ CD3+ CD8a+ CD197- CD45RA+ |

## References

Becht,E. et al. (2018) Dimensionality reduction for visualizing single-cell data using UMAP. Nat. Biotechnol., 37, 38–44.

Bruggner,R.V. et al. (2014) Automated identification of stratifying signatures in cellular subpopulations. Proc Natl Acad Sci USA, 111, E2770–7.

Burton,R.J. et al. (2021) CytoPy: an autonomous cytometry analysis framework. PloS Computational Biology

Luecken,M.D. et al. (2022) Benchmarking atlas-level data integration in single-cell genomics. Nat. Methods, 19, 41–50.

McInnes,L. and Healy,J. (2018) UMAP: Uniform Manifold Approximation and Projection for Dimension Reduction. arXiv [stat.ML].

Oetjen,K.A. et al. (2018) Human bone marrow assessment by single-cell RNA sequencing, mass cytometry, and flow cytometry. JCI Insight, 3.

Pezzotti,N. et al. (2016) Hierarchical Stochastic Neighbor Embedding. Computer Graphics Forum, 35, 21–30.

Qiu,P. et al. (2011) Extracting a cellular hierarchy from high-dimensional cytometry data with SPADE. Nat. Biotechnol., 29, 886–891.

Shahi,P. et al. (2017) Abseq: Ultrahigh-throughput single cell protein profiling with droplet microfluidic barcoding. Sci. Rep., 7, 44447.

Stoeckius,M. et al. (2017) Simultaneous epitope and transcriptome measurement in single cells. Nat. Methods, 14, 865–868.

Sun,S. and Wolf,A. (2022) readfcs: Read FCS files. LnReps.

Traag,V.A. et al. (2019) From Louvain to Leiden: guaranteeing well-connected communities. Sci. Rep., 9, 5233.

Virshup,I. et al. (2021) anndata: Annotated data. BioRxiv.

White,S. et al. (2021) Flowkit: A python toolkit for integrated manual and automated cytometry analysis workflows. Front. Immunol., 12, 768541.

Wolf,F.A. et al. (2018) SCANPY: large-scale single-cell gene expression data analysis. Genome Biol., 19, 15.

